# *maoP* impacts biofilm formation via *csgD* in *Salmonella* Typhimurium

**DOI:** 10.64898/2026.11.20.767877

**Authors:** Joshua C. I. Horton, Emma R. Holden, Eleftheria Trampari, Isabelle Bangs, Jacob G. Malone, Mark A. Webber

## Abstract

Biofilm formation is a key process in the life cycle of most bacteria and involves the formation of multicellular aggregates, frequently attached to a surface. Despite extensive research into the complex regulatory pathways controlling this process, many aspects are not fully understood.

Originally named for its role in chromosomal organisation, MaoP has previously been shown to have an important impact on biofilm formation in *Escherichia coli* and *Salmonella enterica serovar* Typhimurium. Here we continued to explore how MaoP impacts biofilm formation in *S*. Typhimurium 14028*S*. We show that deletion of *maoP* significantly reduces biofilm formation through reduced production of the biofilm matrix components curli and cellulose, and that this is associated with a reduction in transcript abundance of the central biofilm regulator *csgD*. Comparison of the transcriptomes of WT and *maoP* deletion strain biofilms confirmed the substantial impact of MaoP on *csgD* and its regulon.

To further explore how MaoP impacts biofilm formation we conducted a biofilm evolution experiment to select for suppressor mutants that had restored curli production in a Δ*maoP* background. Sequencing of these mutants identified single nucleotide polymorphisms upstream of *csgD* that increased *csgD* transcript abundance and recovered biofilm formation to wild-type levels.

In conclusion, our findings suggest a pathway where MaoP is involved in the regulation of biofilm formation through the control of the central biofilm regulator CsgD.

## INTRODUCTION

Most bacteria form biofilms, aggregates of cells encased within extracellular polymeric substances (EPS) that are often attached to a surface (1). This structure confers protection against various stresses in the external environment. Therefore, bacteria within a biofilm often show a higher tolerance to antimicrobials compared to their planktonic counterparts, meaning that biofilms are often harder to treat via conventional means (2,3). Conversely, biofilms also show potential for use in bioremediation and processing of pollutants such as hydrocarbons, heavy metals, and plastics (4–7). Furthering understanding of the complex regulatory pathways involved in biofilm formation is therefore important to control or exploit this process.

Formation of biofilm is a metabolically expensive process that results in 5% to 50% of all genes demonstrating altered expression in a biofilm compared to planktonic growth (8–11). To facilitate such a dramatic shift in gene expression, biofilm formation is controlled by numerous global regulators (1,12). Cyclic diguanylate monophosphate (c-di-GMP) is a secondary messenger molecule implicated in the transition from the planktonic to the biofilm lifestyle, where high c-di-GMP levels promote biofilm formation whilst repressing genes associated with planktonic growth, including pathways encoding motility and virulence (13–15). One of the main biofilm regulators controlled by c-di-GMP in *Salmonella spp.* is CsgD, which itself regulates the production of key biofilm components, the amyloid fibril protein curli and the polysaccharide cellulose (12,16–18).

Previous work in our group employed transposon mutagenesis screening to discover novel genes important for biofilm formation in *E. coli* and *S. enterica serovar* Typhimurium (*S.* Typhimurium) (19,20). This screen identified *maoP*, encoding a predicted 112 amino acid protein that lacks homology to any previously characterised proteins (21). Named for a proposed role in chromosomal organisation, *maoP* has also been implicated in biofilm-related phenotypes through transposon mutagenesis screens in other Enterobacteriaceae; interruption of *maoP* is associated with increased motility (22) but negatively impacts biotic adhesion (23) and survival in low moisture conditions (24). However, the mechanism by which MaoP influences these processes is yet to be described.

In this study, we investigated the importance of *maoP* in *S.* Typhimurium biofilm formation. Deleting *maoP* reduces biofilm production by decreasing *csgD* mRNA levels, which subsequently reduces the production of the key biofilm components curli and cellulose. Loss of *maoP* also led to significant changes in the transcriptomes of a biofilm, with *csgD* and curli biosynthesis genes markedly repressed in the mutant compared to wild-type (WT). Finally, a laboratory biofilm evolution model demonstrated that Δ*maoP* mutants can evolve to restore normal biofilm formation. The evolved mutants accumulated single nucleotide polymorphisms (SNPs) in the regulatory regions of *csgD*, resulting in increased expression of the master regulator. Collectively these results suggest *maoP* impacts biofilm by promoting the abundance of *csgD*.

## METHODS

### Strains, plasmids, and primers

Bacterial strains used in this study are listed in Table 1. Plasmids and primers used can be found in Supplementary Tables 1 and 2.

**Table 1.** Bacterial strains used in this work.

| Name | Description | Source |
|---|---|---|
| *Salmonella enterica* subsp. *enterica serovar* Typhimurium strain 14028*S* | WT strain | ATCC 14028 |
| WT:: *lacIZ* | LacIZ-tagged WT strain for adhesion competition assay | (25) |
| Δ*maoP* | *maoP* deletion strain | (20) |
| Δ*maoP* + *maoP* | Δ*maoP* with *maoP* under its native promoter inserted at the Tn7 site | This study |

### *maoP* complementation by gene insertion

To complement the Δ*maoP* phenotype, *maoP* under its native promoter was inserted using Gene Doctoring (26) into the chromosome at the Tn7 insertion site using a previously described plasmid (25) (Supplementary Table 1). We have previously shown insertions at this site are phenotypically neutral in biofilm formation (25). The antibiotic resistance marker was removed in the deletion strain prior to gene insertion using pCP20 (27). Plasmid construction was validated by Sanger sequencing (Eurofins) prior to Gene Doctoring. Genetic manipulation in the chromosome was confirmed by Illumina whole genome sequencing.

### Biofilm phenotype assays

Strains were assayed for their biofilm formation ability relative to WT. Biofilm biomass was measured by Crystal violet staining. 200 μl of culture normalised to 10^5^ CFU/ml was added to a polystyrene U-bottom 96-well plate (Grenier bio-one). Plates were sealed with a paper gas-permeable seal (Starlab) and incubated statically for 48 hrs at 30 °C. Planktonic culture was removed by pipetting, plates were gently rinsed with water and left to dry, and then 200 μl 0.1% Crystal violet was added for 10 minutes. The dye was then removed by pipetting, unbound dye rinsed away, and the plates left to dry. The bound Crystal violet was then solubilised by the addition of 200 μl 70% ethanol. Absorbance was measured at 590 nm with a FLUOstar Omega plate reader (BMG Labtech).

Curli and cellulose production were assessed by spotting 5 μl of culture normalised to 10^5^ CFU/ml on LB without salt (LB-NaCl) agar, supplemented with 40 μg/ml Congo red dye (Sigma) or 200 μg/ml Calcofluor White Stain (VWR) for curli or cellulose respectively. Plates were incubated for 48 hours at 30 °C. Congo red plates were photographed. For cellulose production, colonies were resuspended in 2 ml ultrapure water, and 200 μl suspension was added to a 96-well plate. OD_600 nm_ and fluorescence were measured (excitation 360 nm, emission 460 nm) in a FLUOstar Omega plate reader (BMG Labtech). Cellulose production was quantified as a function of Calcofluor fluorescence normalised by OD_600 nm_.

### RNA extraction

RNA was extracted for use in reverse transcription quantitative polymerase chain reactions (RT-qPCR) and RNA-seq to assess transcript abundance. For RNA in a biofilm, 5 μl of culture at 10^5^ CFU/ml was spotted on LB-NaCl agar and grown for 24 hours at 30 °C. The colony was resuspended in 1 ml PBS in a microcentrifuge tube, pelleted by centrifugation at max speed for 1 minute, and the supernatant removed. For mRNA levels at mid-exponential growth, 10 ml LB-NaCl broth was inoculated with 10^7^ CFU/ml culture and grown to an OD_600 nm_ of 0.3-0.4, at 30 °C 180rpm. The culture was then centrifuged at 3000 x *g* for 10 minutes and most of the supernatant poured off. The pellet was resuspended in the remaining supernatant, transferred to a microcentrifuge tube and pelleted by centrifugation at max speed for 1 minute and the remaining supernatant removed.

RNA was then extracted for either growth condition using the Quick-RNA™ Fungal/Bacterial Miniprep Kit (Zymo Research) as per manufacturer’s instructions, including the on-column DNase digestion using DNase I Set (Zymo Research). RNA was stored at −20 °C for short term, and −70 °C for long term storage.

RNA quantity and purity was measured using Qubit High Sensitivity RNA quantification (Invitrogen) and NanoDrop, with DNA contamination being checked for with QuantiFluor dsDNA kit (Promega). RNA quality was assessed using an Agilent 2200 TapeStation with a High Sensitivity RNA ScreenTape as per manufacturer’s protocol. Only samples with an RNA Integrity Number (RIN) greater than 5 were used for RT-qPCR and 6 for RNA-seq. RNA samples for RT-qPCR were further processed to remove contaminating DNA using the Turbo DNA-free™ kit (Invitrogen).

### Reverse transcription quantitative polymerase chain reaction (RT-qPCR)

mRNA levels of target genes were measured by RT-qPCR. Primers were designed using Primer3 (28) and checking for dimers or secondary structures using the IDT Oligo Analyser tool. Primer amplification efficiency was validated against genomic DNA extracted from WT.

RT-qPCR reactions were carried out in 10 μl volumes using Luna® One-Step Universal RT-qPCR kit (NEB) as per the manufacturer’s protocol. For each sample and for each target gene, controls with no reverse transcriptase were included.

mRNA levels were calculated as fold-change relative to the housekeeping gene *gyrB* using the 2^ΔΔCt^ method (29).

### RNA-seq

RNA samples were sequenced by Genewiz (Azenta Life Sciences, UK): strand-specific RNA-sequencing after rRNA depletion produced approximately 10 million reads per sample. Three biological replicates for each condition were sequenced.

Data were analysed using a previously described analysis pipeline (30). Reads were quality control trimmed by fastp v0.23.2 (31) and aligned to the reference genome of *Salmonella* Typhimurium 14028*S* (GenBank accession number GCA_000022165.1) using HISAT2 v2.10 (32,33). Featurecounts v1.6.3 (34) was used to assign aligned reads to annotated features, altering the feature type parameter to ‘CDS’ or ‘transcript’, and the GFF gene identifier to ‘locus_tag’, before DESeq2 v2.11.40.8 (35) was used to determine differentially expressed genes. The output file was then processed by Annotate DESeq2/DEXSeq output tables v1.1.0 with the gff feature type parameter being ‘CDS’ or ‘transcript’, GFF feature identifier parameter was set as ‘locus_tag’, and GFF attributes to include parameter was set to ‘gene’ and ‘product’.

The generated files were then merged and processed with R (version 4.6.0).

### Biofilm evolution model

To investigate the mechanisms of increased biofilm formation in Δ*maoP*, we used a previously described model that selects for increased biofilm biomass production (36). Δ*maoP* and WT biofilms were grown on alternating 6 mm and 3 mm diameter stainless steel beads (Simply Bearings Ltd.) in 5 ml LB-NaCl broth and incubated at 30 °C with gentle shaking at 80 rpm. 3 lineages of each strain were set up by inoculating with 10^7^ CFU/ml of overnight culture. Cultures were passaged every 48-72 hours by taking one bead, washing briefly in sterile PBS, and transferring to fresh media with sterile beads of the other size (allowing the ‘incoming’ and new beads to be distinguished).

A sample of the model was taken every 3^rd^ passage: a bead was briefly washed in sterile PBS to remove planktonic cells and then vortexed vigorously in 500 μl PBS to resuspend the biofilm. 50 μl of resuspended biofilm in PBS was then used to inoculate 5 ml LB and grown overnight at 37 °C 200 rpm. A sample was then stored in 10% glycerol at −20 °C.

Isolates were recovered from the biofilm populations in PBS by serial dilution and spreading on LB-NaCl agar supplemented with Congo red dye (40 μg/ml). Colonies showing increased curli production, either partial or full recovery to WT levels, were selected and used to inoculate an overnight culture. These were subsequently stored in 10% glycerol at −20 °C for phenotyping and whole genome sequencing. Genomic DNA was extracted as described previously (36) and eluted into Nuclease-Free Water (Sigma). Short-read libraries, of DNA normalised to 5 ng/μl, were prepared as previously described (37) using a NextSeq™ 2000 P4 XLEAP-SBS™ Reagent Kit (300 Cycles) (Illumina). The libraries were pooled following quantification in equal quantities, and the pool run at a final concentration of 750 pM on an Illumina Nextseq 2000.

### SNP analysis

Illumina reads of evolved isolates were run through Snippy4 (version 4.4.3) (https://github.com/tseemann/snippy), comparing against the reference genome for the parental strain. The ‘minimum proportion for variant evidence’ option was reduced to 0.1, from the default 0.9, to ensure all SNPs are identified, even at low occurrence. A selection of SNPs was verified by PCR and Sanger sequencing (Eurofins).

The outputs were merged into one datasheet, synonymous SNPs removed, and then manually reviewed to exclude likely sequencing errors (e.g. in repeat regions). The output was manually checked against the reference genome to determine the upstream region descriptions, noting the position relative to the predicted start codon.

### Biofilm adhesion competition assay

Adhesion by mutant strains was measured in competition with a LacIZ tagged WT strain that has previously been shown to be phenotypically neutral in growth and competition assays (25,38), as well as in this screen (Supplementary Figure 2). This tag allowed for differentiation between mutant and WT populations by blue-white screening. Sterile 6 mm stainless steel beads (Simply Bearings Ltd.) were added to universal tubes with 5 ml LB-NaCl broth. Overnight culture of the tagged WT and mutant strains were normalised to the lowest OD_600 nm_ before 50 μl of each culture of tagged WT and mutant strain was added to the tube with beads and incubated at 30 °C with mild agitation at 80 rpm.

CFU percentage was determined by serial dilution and spotting on LB-NaCl agar plates supplemented with X-gal (40 μg/ml) and 1 mM isopropyl β-D-1-thiogalactopyranoside (IPTG). At 0 hours, a 20 μl sample of the LB-NaCl broth was taken for serial dilution and CFU counts. After 3, 6, and 12 hours, one stainless steel bead was removed, washed briefly in PBS, before being placed into 500 μl fresh PBS in a microcentrifuge tube. The tube was then vortexed vigorously for 10 seconds, and 20 μl of PBS was taken for serial dilution for CFU counts. Plates for CFU counts were incubated overnight at 37 °C.

## RESULTS

### Deletion of *maoP* reduces biofilm formation through *csgD*

Previously we showed that deletion of *maoP* in *S.* Typhimurium resulted in a reduction in biofilm biomass production when compared to WT (20). To further investigate the role that *maoP* has in biofilm formation, we tested how deletion of *mao*P affects biomass production, curli and cellulose synthesis and whether these phenotypes could be complemented by reintroducing *mao*P in the chromosome.

A complementation strain was created by inserting *maoP* with its native promoter at the Tn7 insertion site adjacent to *glmS* (25). We found that upon reinsertion of *maoP*, biofilm biomass production was restored levels significantly equivalent to WT (TOST, *p* < 0.001) (Figure 1a).

**Figure 1.**
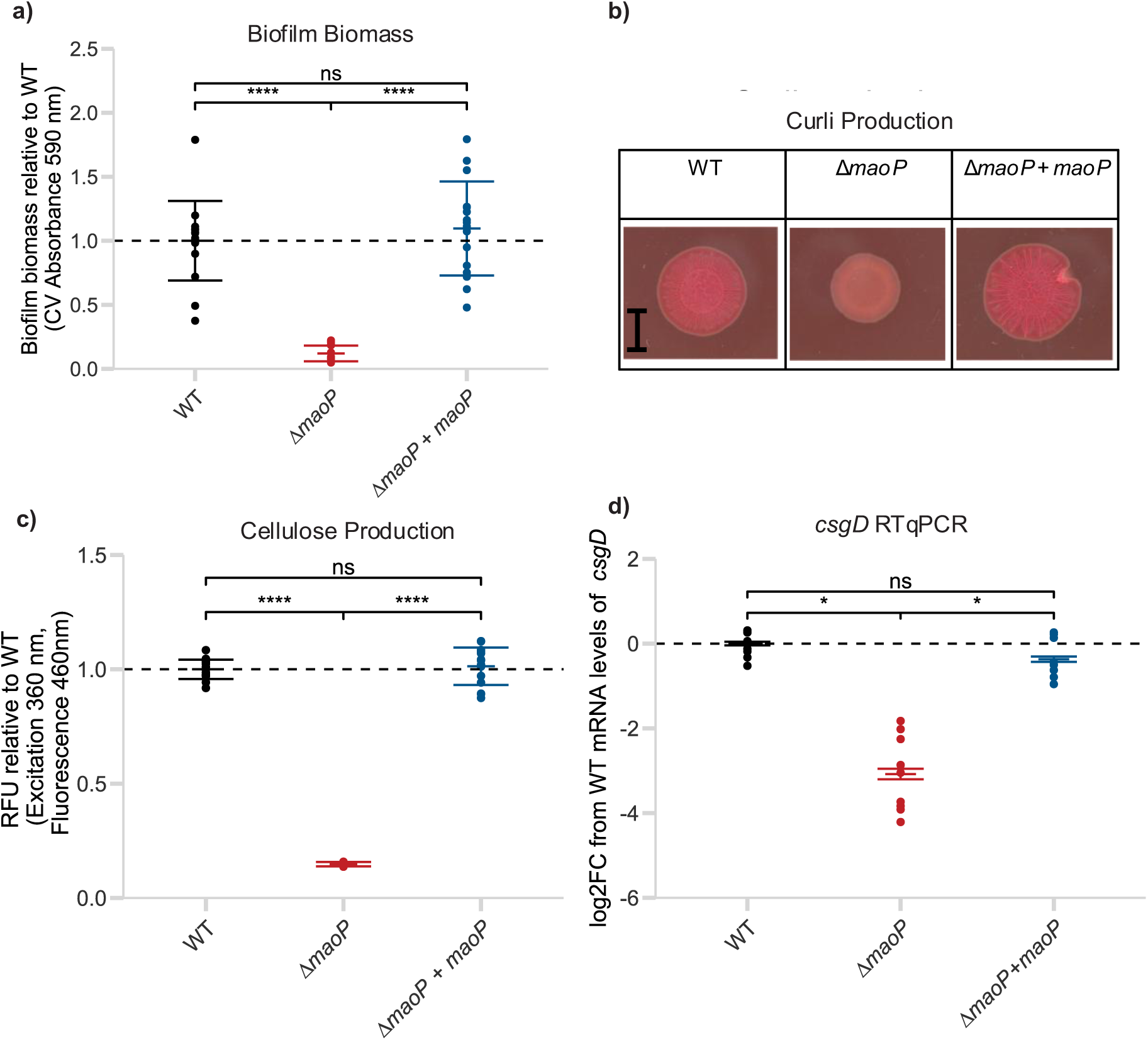
Deletion of *maoP* reduces biofilm formation due to reduced *csgD* mRNA in the cell. **(a)** Biofilm biomass, relative to WT, measured by Crystal violet (CV) staining. Points show 16 replicates (eight samples from each of two independent runs completed on different days). **(b)** Curli production visualised by Congo red staining. Images are representative of six replicates (three samples from each of two independent runs completed on different days). Scale bar = 0.5 cm. **(c)** Cellulose production, relative to WT, measured by Calcofluor staining. Points show 16 replicates (eight samples from each of two independent runs completed on different days). Error bars in **(a)** & **(c)** show one standard deviation. Statistical significance was calculated by one-way ANOVA with Tukey’s post-hoc analysis (****: *p*<0.0001; *: *p*<0.05 ns: *p*>0.05). **(d)** Log_2_-fold change (log2FC) in *csgD* mRNA abundance relative to WT, relative to *gyrB* as a housekeeping gene. RNA extracted from a 24-hour biofilm plated on LB without NaCl agar. Error bars around the mean show standard error of the mean (SEM). Points shown nine replicates from three independent experiments of three replicates each. Statistical significance was calculated on ΔCt values by one-way ANOVA and Tukey’s post-hoc analysis (* : *p*<0.05; ns: *p*>0.05).

To further characterise the biofilm-deficient phenotype, we investigated the production of two key biofilm matrix components; curli and cellulose. Curli is an amyloid fibre protein that plays a key role in biofilm formation in *Salmonella spp.* (39). Production of curli can be visualised by growth on agar supplemented with Congo red dye (40). WT shows a ‘red, dry, and rough’ (rdar) phenotype, indicating high curli production. Upon deletion of *maoP*, we found that *S.* Typhimurium produces a pink smooth colony, consistent with reduced curli production, which recovers to the rdar phenotype upon complementation of *maoP* (Figure 1b). Cellulose is another key component of biofilm in many Enterobacteriaceae (41). Cellulose production can be measured by fluorescence of the Calcofluor White stain that binds to cellulose (42). Deletion of *maoP* resulted in a significant reduction in cellulose production compared to WT, which was restored in the complementation strain (Figure 1c).

Often described as the central regulator of biofilm in *S.* Typhimurium, CsgD controls among others the production of both curli (directly) and cellulose (indirectly) (43). To determine if the changes in matrix production were due to changes in *csgD* mRNA levels in the cell, RT-qPCR was used. Δ*maoP* showed an 8-fold reduction in *csgD* mRNA levels in a biofilm compared to the WT strain, which was successfully complemented by reinsertion of *maoP* in the chromosome (Figure 1d).

Taken together, these data show that loss of *maoP* results in decreased biofilm formation, likely due to a reduction in *csgD* mRNA levels causing a drop in the production of key biofilm components curli and cellulose.

### Impact of loss of *maoP* on the biofilm transcriptome

To understand the global impact on the transcriptome caused by an absence of *maoP* in a biofilm, RNA-seq was used. Using a previously described analysis pipeline (30), we compared the transcriptome of WT and Δ*maoP* strains in a 24-hour biofilm. We found that deletion of *maoP* resulted in 8% of annotated genes in the reference genome showing a log_2_ fold-change of greater than 1 or less than −1, and an adjusted *p*-value of less than 0.05 (452 out of 5591 CDS and transcripts) (Figure 2).

**Figure 2.**
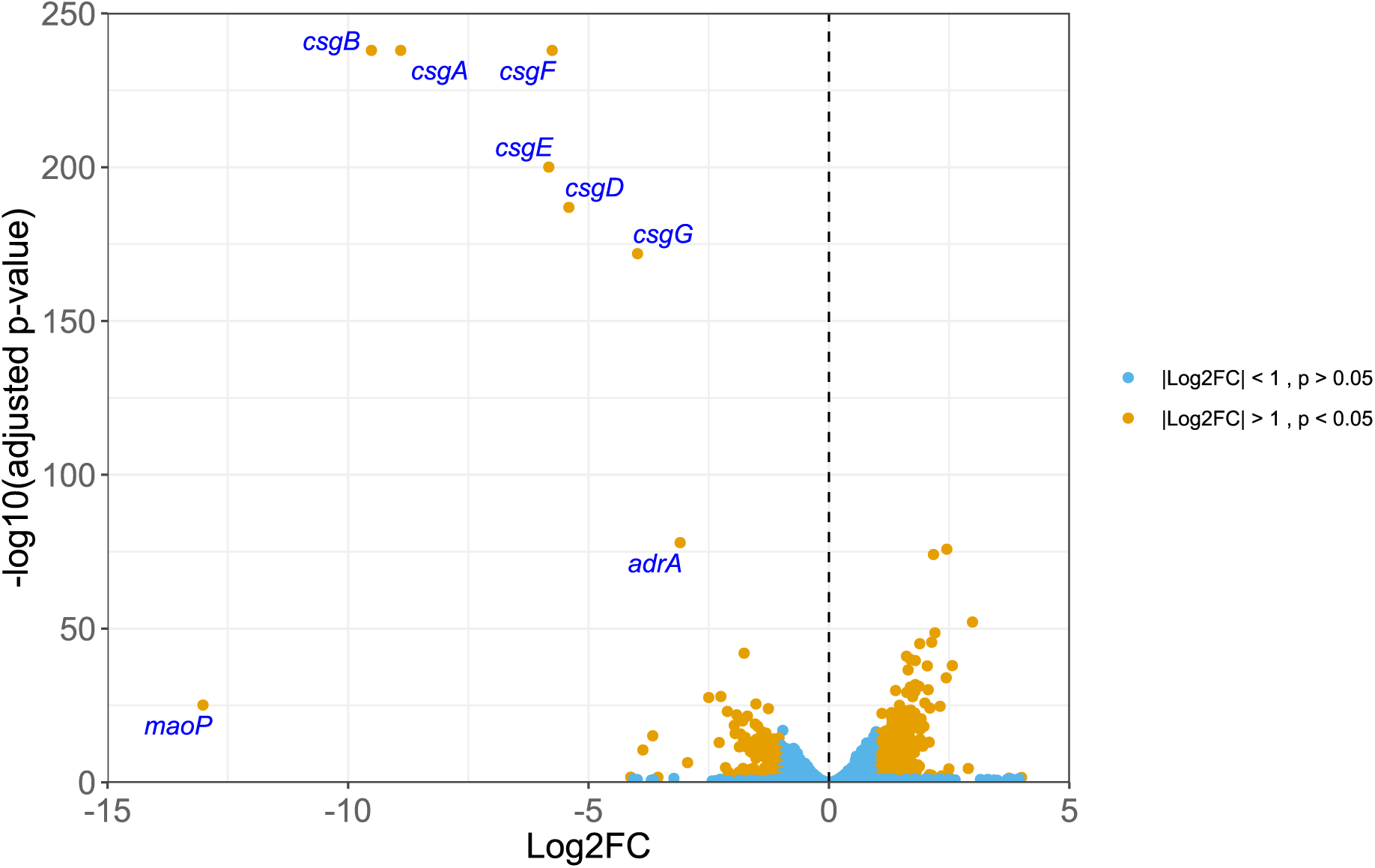
Deletion of *maoP* has a global effect on the transcriptome of a 24-hour biofilm, especially on key biofilm genes. X-axis displays log_2_-fold change (Log2FC) in gene expression, where a positive value indicates upregulation in Δ*maoP* relative to WT, and a negative value indicates downregulation. Y-axis represents statistical significance as the -log_10_ adjusted *p*-value calculated by DESeq2. Dashed line indicates no change in expression (x=0). Colour denotes significance. Genes with the seven lowest adjusted *p*-values are labelled, with the addition of *maoP*.

To identify networks of genes differentially expressed in the Δ*maoP* 24-hour biofilm compared to WT 24-hour biofilm, we submitted the differentially expressed genes to the STRING database (44). We found that genes involved in flagellar assembly and chemotaxis (STRING cluster CL:5089, 48 out of 52 in the network), as well as virulence (STRING cluster CL:5324, 31 out of 46 in the network) were found to be differentially expressed, in addition to those known to be important for biofilm, and various metabolic pathways (Supplementary Figure 1).

Of the genes with the lowest adjusted *p*-value and largest differential expression, six out of seven genes from the two *csg* operons (except *csgC*), involved in biofilm regulation and curli production including *csgD* were strongly down-regulated (16,17). In addition, the diguanylate cyclase *adrA*, whose expression is itself regulated by CsgD and activates cellulose production, was also found to be within the seven annotated genes with the lowest adjusted p-value (45–48). This shows that the most prominent effect that deletion of *maoP* has on the transcriptome is on key genes involved in the regulation of biofilm formation with the curli biosynthesis and regulation pathway being the most significantly affected. However, the way in which MaoP causes this effect is unclear.

### Selection for increased biofilm formation can rescue the *maoP* deficit but the trajectory differs in the presence and absence of *maoP*

To try to uncover how MaoP impacts biofilm formation in an unbiased way, we selected for mutants where the biofilm deficit caused by a loss of *maoP* was alleviated. A previously optimised biofilm evolution model was used to select for increased biofilm formation in both WT and Δ*maoP* by repeated passage on stainless-steel beads (36,49) (summarised in Figure 3a). This model has been previously shown to select for increased biofilm formation over time (36,49). Isolates of Δ*maoP* with increased Congo red retention (suggestive of increased curli production) were recovered after seven passages. This was longer than the four passages required for increased curli production to be selected for in the WT. Representative colonies of those that demonstrated either intermediate recovery (darker pink/light red colonies), or full recovery of curli production (rdar) in the Δ*maoP* background were isolated throughout the 15 passages used in the model and subjected to whole genome sequencing. Colonies were also recovered from WT lineages on the basis of increased Congo red dye retention and sequenced.

**Figure 3.**
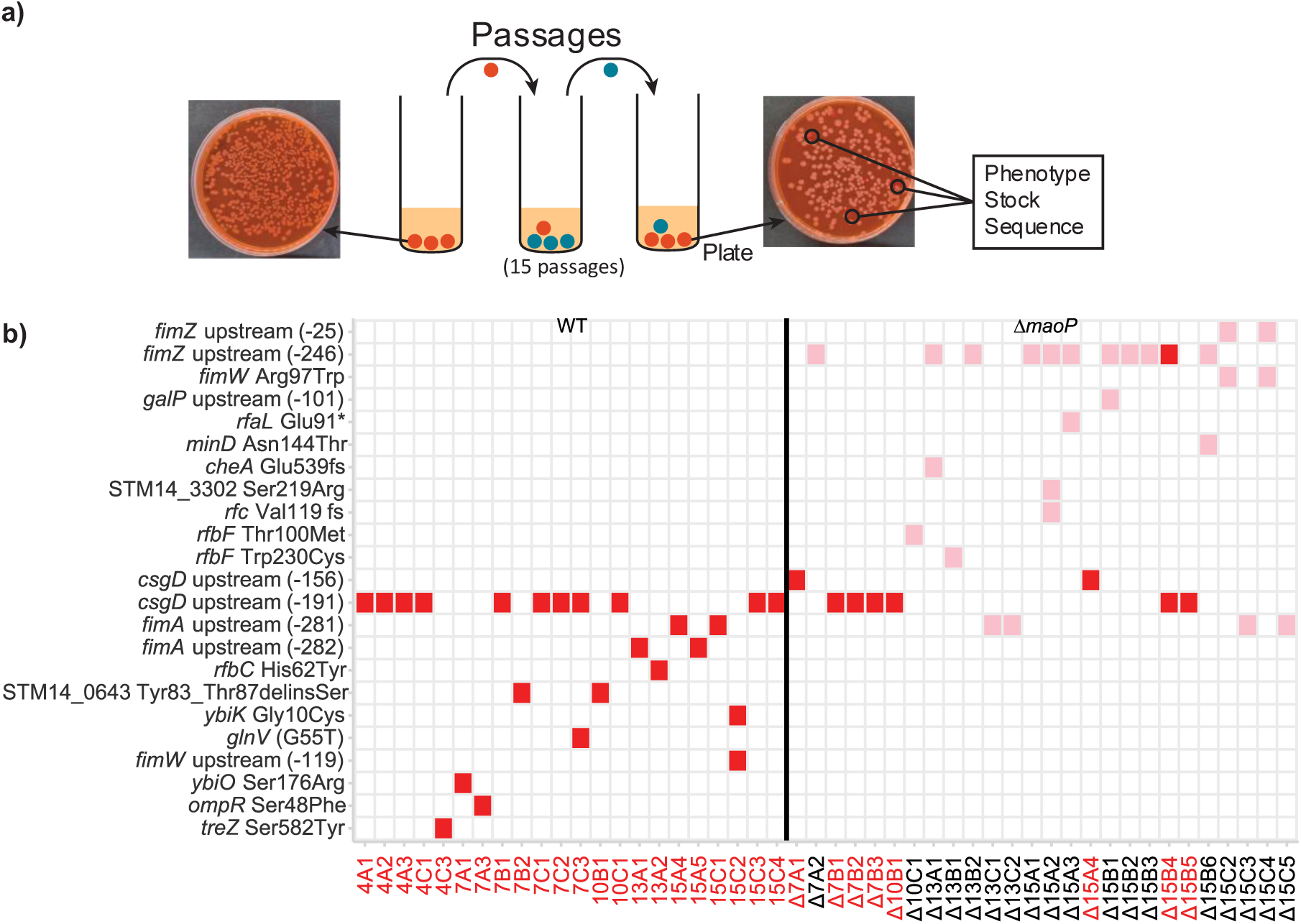
*S.* Typhimurium evolves differently in the presence and absence of *maoP* in a biofilm evolution model. **(a)** Schematic of the biofilm evolution model used. **(b)** Single nucleotide polymorphisms (SNPs) identified in WT and Δ*maoP* mutants selected in the biofilm evolution model. Numbers in brackets denote bases upstream from the start codon. Fs= frameshift. Delins= deletion/insertion. * = premature stop codon. Isolate names dictate passage number, lineage, and isolate number, respectively. Red squares/text indicate strains that presented the rdar phenotype on Congo red plates. Pink squares indicate strains with an intermediate phenotype.

Analysis of the mutations present in the recovered colonies showed a distinct evolutionary trajectory in the presence and absence of *maoP* (Figure 3b). Only two SNPs were shared in both WT and Δ*maoP* isolates recovered from the model. These were both likely regulatory changes present upstream of *csgD* (191 bases upstream from the start codon) or upstream of *fimA* (281 bases upstream from the start codon).

### Changes upstream of *csgD* rescue the biofilm defect of Δ*maoP* mutants and increase *csgD* mRNA abundance

Δ*maoP* isolates were recovered from the model that had reversed the biofilm-deficient phenotype of the parent strain and recovered the rdar phenotype on Congo red plates. Upon whole genome sequencing, all such isolates possessed one of two SNPs upstream of *csgD*. Also found in WT evolved isolates, one SNP was 191 bases upstream of the start codon and was located in the OmpR binding site. Part of the EnvZ-OmpR two-component system that responds to environmental osmolarity, OmpR activates *csgD* transcription when not phosphorylated, at which point it represses expression, with the mutated binding site being from where OmpR promotes *csgD* transcription (43,50,51). The other SNP, found solely in evolved Δ*maoP* isolates, was 156 bases upstream of the start codon, which is predicted to lie in the region of the *csgD* promoter that directs RNA polymerase binding. (Figure 4a).

**Figure 4.**
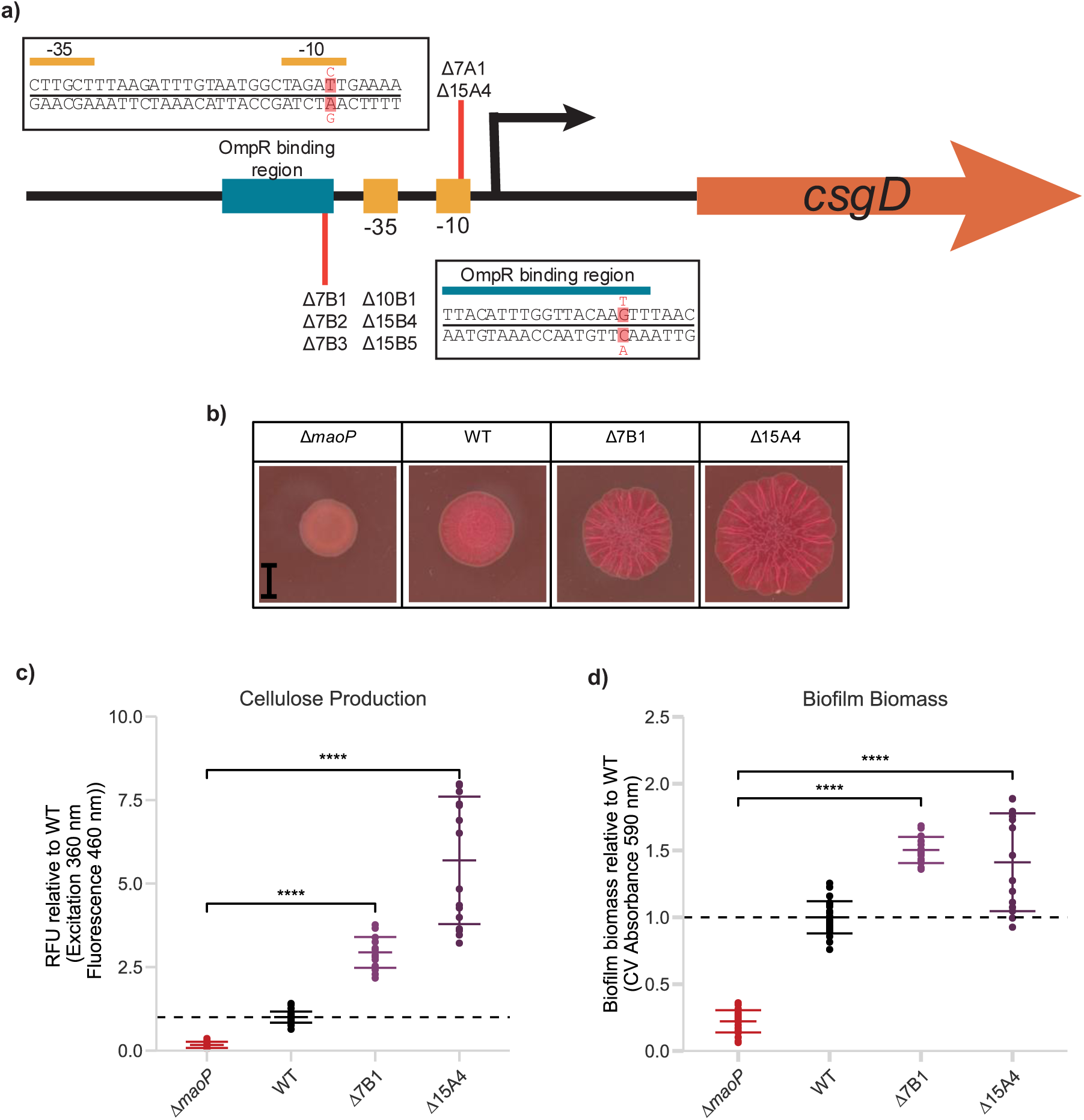
Δ*maoP* evolves SNPs upstream of *csgD* in a biofilm evolution model that significantly increases biofilm formation. **(a)** Location of SNPs identified upstream of *csgD*. **(b)** Curli production visualised by Congo red dye staining. Images are representative of six replicates (three samples from each of two independent runs completed on different days). Scale bar = 0.5 cm **(c)** Cellulose production, relative to WT, measured by Calcofluor staining. Points show 16 replicates (eight samples from each of two independent runs completed on different days). **(d)** Biofilm biomass production, relative to WT, measured by CV staining. Points show 16 replicates (eight samples from each of two independent runs completed on different days). Error bars in **(c)** & **(d)** show one standard deviation. Statistical significance was calculated by one-way ANOVA with Tukey’s post-hoc analysis (****: *p*<0.0001)

Δ7B1 and Δ15A4 were chosen as representative strains with one of the SNPs upstream *csgD* to carry forward for further characterisation as these did not possess any other SNP. Δ7B1 has the SNP located in the OmpR binding region (upstream −191), whilst Δ15A4 has the SNP located in the −10 box (upstream −156).

Both isolates demonstrated significantly increased biofilm formation, including visibly increased curli and significantly increased cellulose production when compared to the parent Δ*maoP* strain (Figure 4).

We hypothesised that this increase in biofilm formation was due to an increase in *csgD* mRNA levels in the cell, compensating for the decrease found in the Δ*maoP* parent strain. RT-qPCR confirmed that *csgD* mRNA levels in a biofilm were substantially increased compared to Δ*maoP* in strains with either SNP (Figure 5).

**Figure 5.**
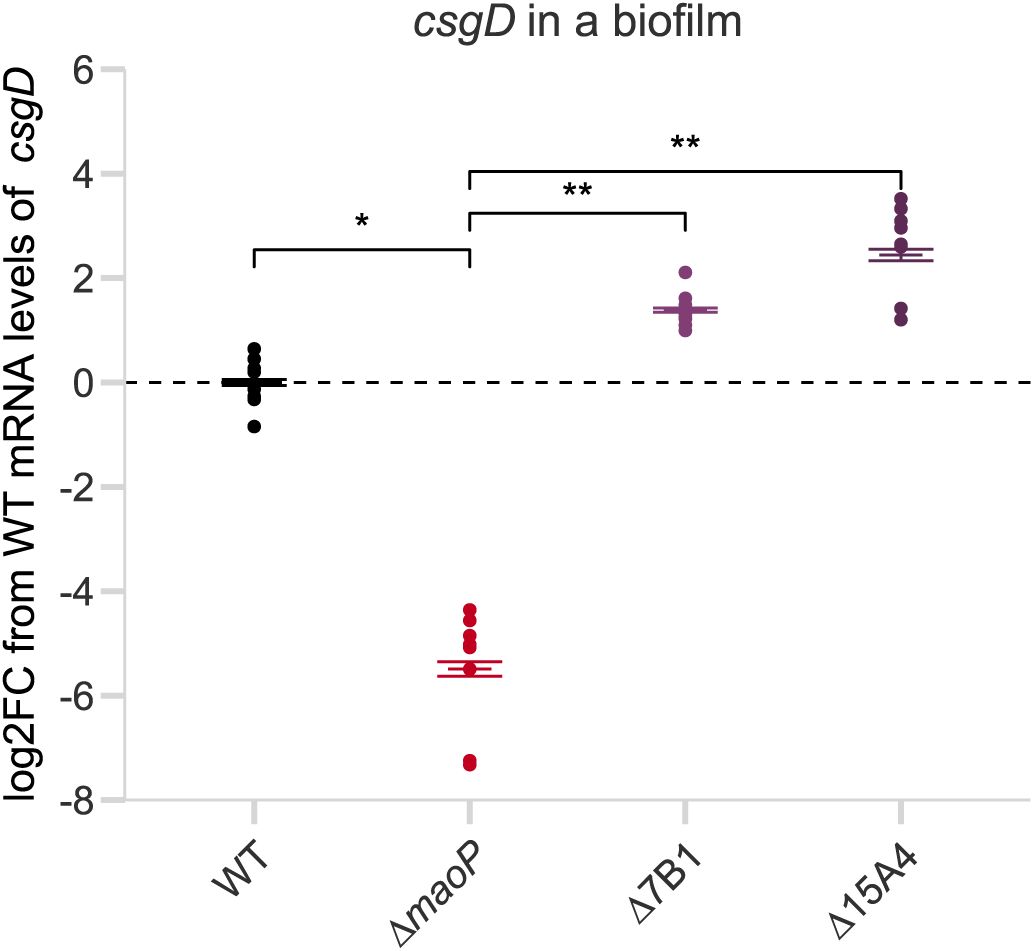
SNPs upstream of *csgD* in Δ*maoP* increase *csgD* mRNA levels. Log_2_-fold change (log2FC) in *csgD* mRNA levels measured by RT-qPCR, relative to WT and relative to *gyrB* as a housekeeping gene. RNA was extracted from a 24-hour biofilm grown at 30 °C on LB-NaCl agar. Error bars show SEM. Points show nine replicates (representing three samples from each of three independent runs completed on different days). Statistical significance was calculated on ΔCt values with a one-way ANOVA with Tukey’s post-hoc analysis (* : *p*<0.05; ** : *p*<0.01).

### Increased adhesion selects for SNPs upstream of *fimZ* in Δ*maoP* biofilm evolution

Isolates were also observed that showed intermediate recovery on Congo red dye supplemented agar plates (i.e. colonies that were darker pink than surrounding colonies but not rdar) and a selection of such isolates were recovered as it was hypothesised they could reveal alternative targets of *maoP* with importance in biofilm formation. Half of all Δ*maoP* isolates with SNPs relative to the reference genome (13 out of 26) had changes upstream of the positive regulator of fimbriae production, *fimZ* (Figure 3b). These changes were restricted to the Δ*maoP* background and were not seen in the WT context. Fimbriae are rod-like structures that protrude from the cell and have been shown to be important in the adhesion stages of biofilm formation (8,9,11). More recently they have also been implicated at each stage of biofilm formation (20).

To further investigate the importance of these mutations, representative strains carrying one of the two SNPs (summarised in Figure 6a) were chosen: Δ7A2 with a SNP 246 bases upstream of the start codon of *fimZ* and Δ15C2 with a SNP 25 bases upstream of the *fimZ* start codon. The former is predicted to be within the −10 and −35 binding regions of the RNA polymerase while the latter is predicted to be within the 5’ untranslated region (5’UTR), but before the Shine-Dalgarno sequence. The SNP 25 bases upstream of the *fimZ* start codon only cooccurred with a SNP within the coding region of *fimW*, encoding the negative regulator of fimbriae. However, this SNP was not predicted to be within the helix-turn-helix domain thought to be important in its role of negatively regulating *fimY* expression (52).

**Figure 6.**
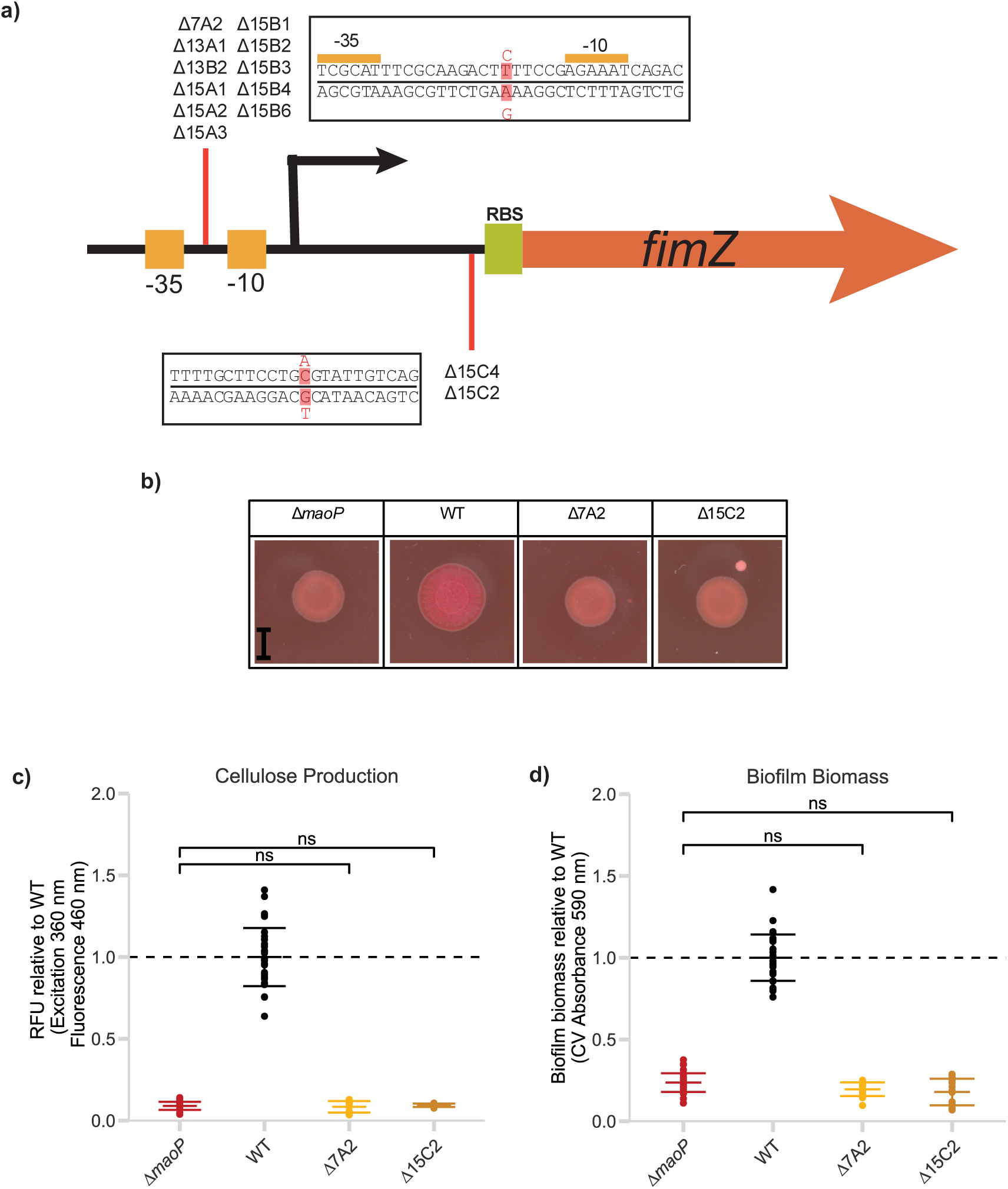
*maoP* deletion strain with SNPs upstream of *fimZ*, selected for in a biofilm evolution model, do not show altered biofilm formation to the parent strain. **(a)** Location of SNPs identified. RBS = Ribosome binding site. **(b)** Curli production visualised by Congo red dye staining. Images are representative of six replicates (three samples from each of two independent runs completed on different days). Scale bar = 0.5 cm **(c)** Cellulose production, relative to WT, measured by Calcofluor staining. Points show 16 replicates (eight samples from each of two independent runs completed on different days). **(d)** Biofilm biomass production, relative to WT, measured by CV staining. Points show 16 replicates (eight samples from each of two independent runs completed on different days). Error bars in **(c)** & **(d)** show one standard deviation. Statistical significance was calculated by one-way ANOVA (ns : *p*>0.05).

These strains were subjected to biofilm phenotyping as before. Despite being selected as isolates with intermediate recovery, neither strain showed a significant increase in curli, cellulose, or biofilm formation relative to the parent Δ*maoP* strain (Figure 6).

We hypothesised that these SNPs compensated for a difference in *fimZ* mRNA levels in the Δ*maoP* parent strain. We therefore determined the mRNA levels of *fimZ* in a biofilm and in mid-exponential growth for WT, Δ*maoP*, Δ7A2, and Δ15C2 (Figure 7). We saw a significant increase in *fimZ* mRNA levels in Δ7A2 in a biofilm. However, selection for this change is not likely to be a response to a deficiency in the Δ*maoP* strain as the parent strain showed no statistically significant difference in *fimZ* mRNA levels compared to WT. The same trend was observed at mid-exponential growth.

**Figure 7.**
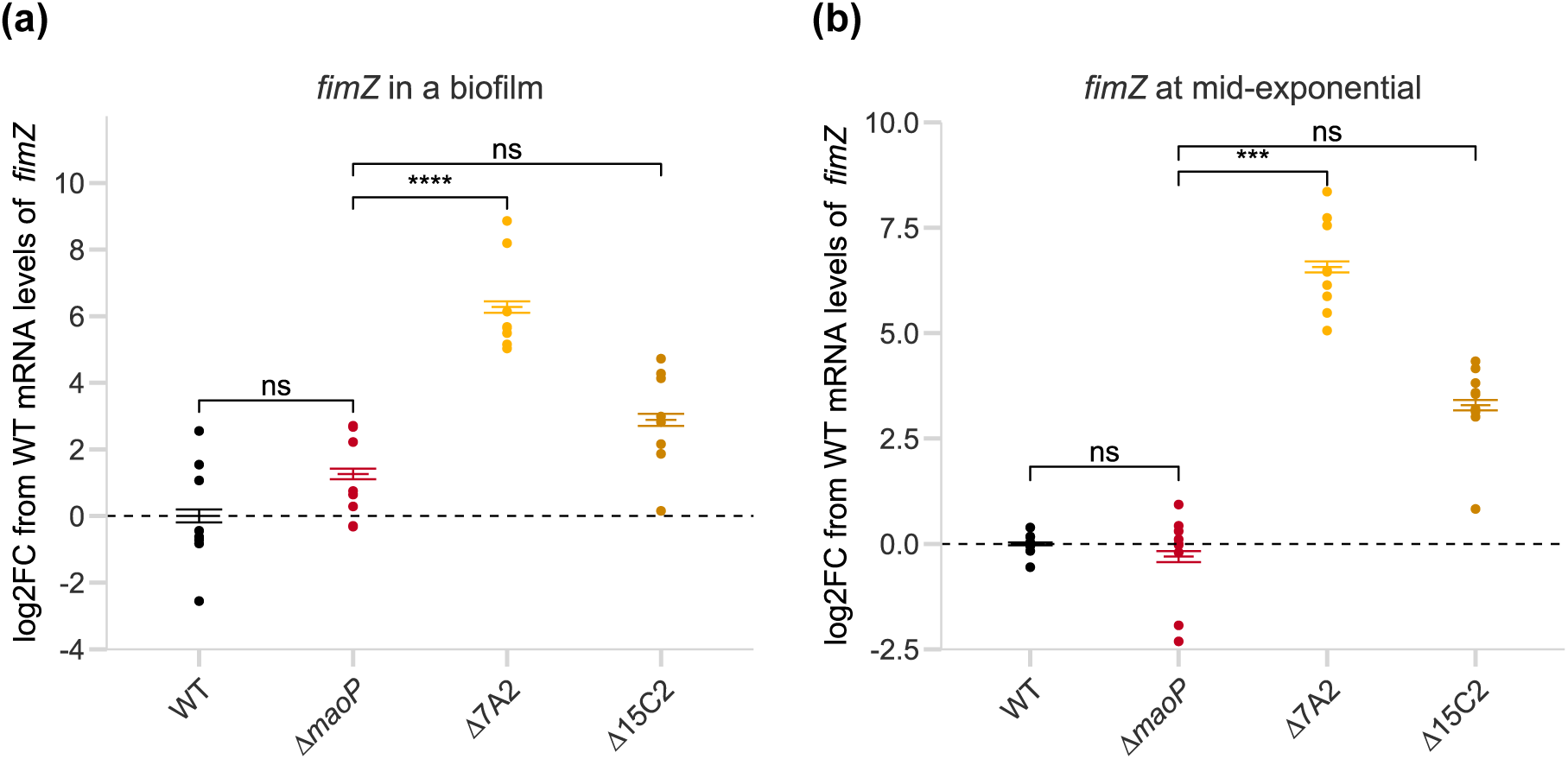
SNPs upstream of *fimZ* increases transcript abundance in both biofilm and mid-exponential in a Δ*maoP* background. Log_2_-fold change (log2FC) in *fimZ* mRNA levels, measured by RT-qPCR, relative to WT and relative to *gyrB* as a housekeeping gene. **(a)** RNA was extracted from a 24-hour biofilm grown at 30 °C on LB-NaCl agar. **(b)** RNA was extracted from a mid-exponential culture in LB-NaCl broth. Error bars show SEM. Points show 9 replicates (representing three samples from each of three independent runs completed on different days). Statistical significance was calculated with ΔCt values using a one-way ANOVA and Tukey’s post-hoc analysis (**** : *p*<0.0001; *** : *p*<0.001; ns : *p*>0.05).

The lack of difference in *fimZ* mRNA levels between WT and Δ*maoP* suggested that mutations were not arising to compensate for a deficiency in transcript abundance of *fimZ* in the Δ*maoP* strain, but instead we hypothesised they provide an alternative route to improved fitness in this biofilm evolution model. As fimbriae are important for adhesion, we hypothesised that although these SNPs did not appear to impact biofilm matrix production, they were selected for in the biofilm evolution model due to increased surface adhesion, and hence increased competitiveness, compared to the parent Δ*maoP* strain.

To test this hypothesis, we measured CFUs of mutant strains bound to stainless-steel beads during the early adhesion to early biofilm stages (3-12 hours) of biofilm formation, in competition with a LacIZ tagged WT strain (25,38) (supplementary Figure 11). We found that the strains harbouring SNPs upstream of *fimZ* showed a statistically significant increase in fitness when competed against the tagged WT for early biofilm when compared to the Δ*maoP* parent strain at 6 hrs (Δ7A2 only) and 12 hrs (both) (Figure 8).

**Figure 8.**
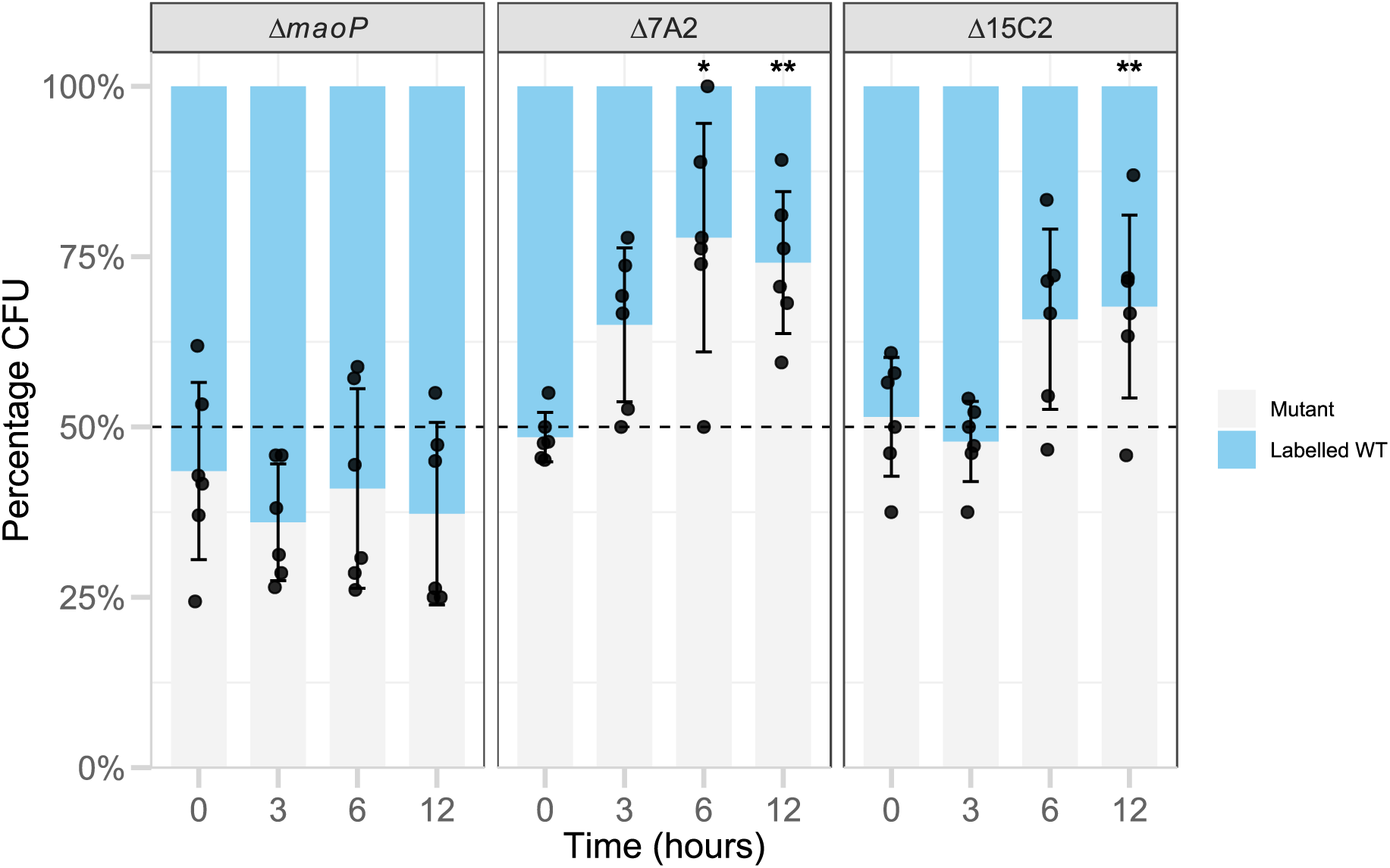
SNPs upstream of *fimZ* in a Δ*maoP* background show increased CFU in early biofilm when in competition with a labelled WT strain. Biofilm adhesion competition assay between a LacIZ tagged WT strain and Δ*maoP* or evolved isolates with SNPs upstream of *fimZ*. CFU counts taken at the Time=0 (planktonic), 3, 6, and 12 hours (biofilm). The mean percentage of mutant (white) and labelled WT (blue) colonies are represented by the grey and blue bars respectively. Error bars show one standard deviation of the percentage of mutant colonies. Points represent six replicates (representing three samples from each of two independent runs completed on different days). Statistical significance between the Δ*maoP* parent strain and the selected mutants at each time point was determined by a one-way ANOVA with Tukey’s post-hoc analysis (* : *p*<0.05; ** : *p*<0.01, unlabelled : *p*>0.05).

These data suggest that these SNPs result in increased fitness in the evolution model conditions due to increased adhesion in the early biofilm. The fact these mutants were only recovered from the Δ*maoP* background may suggest their benefit is limited to this background due to the baseline fitness of WT being higher in the model.

## DISCUSSION

Despite extensive work investigating the regulation of biofilm formation in *S.* Typhimurium, much remains to be understood. Previous work in our group uncovered the importance of *maoP* in biofilm formation (19,20) but without any mechanistic insight as to how it exerts its impact. Here, we build upon this previous work to further investigate the involvement of MaoP in biofilm formation in *S.* Typhimurium.

### Deletion of *maoP* reduces *csgD* levels in the cell, resulting in a biofilm deficient phenotype

We previously found that deletion of *maoP* results in a significant reduction in overall biofilm biomass as well as a visible reduction in curli production (20). Here we show that this extends to significantly reduced cellulose production.

The central regulator of biofilm in *S.* Typhimurium, CsgD, is known to regulate both curli and cellulose production (12,16,17). We therefore investigated the effect that deletion of *maoP* has on *csgD* mRNA levels in biofilm cells, and this showed a statistically significant decrease relative to the WT strain.

RNA-seq analysis on the impact on the biofilm transcriptome caused by deletion of *maoP* also demonstrated a marked reduction in *csgD* transcript abundance, by approximately 11-fold. Furthermore, the seven genes with the lowest adjusted *p*-values were directly regulated by *csgD*, including six of the seven genes in the *csg* operons, and *adrA* (45). *adrA* encodes a diguanylate cyclase which produces the key biofilm secondary molecule c-di-GMP which in turn activates the cellulose production machinery and contributes to the global cell pool of c-di-GMP (47). These results highlight that the most significant impact that deletion of *maoP* has on the cell is the reduction of *csgD* transcript abundance, with the reduced curli, cellulose, and overall biofilm formation phenotypes seen all being consistent with this reduction.

Deletion of *maoP* also had a large global effect on the biofilm transcriptome, including motility and virulence pathways. We hypothesise that the majority of such affected genes are changed in response to the primary impact on *csgD* transcript abundance, which causes alterations in the global cell pool of c-di-GMP as well as metabolism caused by the different conditions within a Δ*maoP* biofilm.

Despite extensive analysis, the RNA-seq data did not reveal a clear connection for how MaoP impacts biofilm formation through *csgD*. There were no transcriptional regulator(s), other than *csgD*, substantially up or downregulated in the Δ*maoP* biofilm, nor was there any signal suggesting that MaoP interacts with one of the many regulators known to impact *csgD*. However, as the *S*. Typhimurium 14028*S* reference genome lacks annotation for all sRNAs known to regulate *csgD*, an interaction with such species is plausible. Given that *E. coli* MaoP has been described as an RNA-binding protein (53–55), yet its specific RNA targets remain unknown, work is ongoing to investigate the RNA species bound by MaoP that lead to reduced *csgD* levels, including whether MaoP directly targets *csgD* mRNA or its regulatory sRNAs.

Alternatively, work in the MaoP homolog in *Yersinia pestis* has demonstrated its ability to bind to and alter the structure of DNA as a nucleoid-associated protein (56). CHIP-seq demonstrated a global binding ability of MaoP in *Y. pestis*, including in its biofilm regulation pathway which is substantially different to that of *S.* Typhimurium. Unfortunately, the authors were unable to identify a binding motif. In addition, MaoP from *Y. pestis* or *E. coli* was able to complement the deletions of the other organism, suggesting a similar mode of action which likely extends to *Salmonella spp*. Therefore, future work will also focus on whether MaoP, in *S.* Typhimurium, binds directly with the *csgD* promoter DNA sequence and promote expression.

### *Salmonella* Typhimurium follows a distinct evolutionary trajectory in the presence or absence of MaoP

To further unpick the complex impact that deletion of *maoP* has on biofilm formation, we used a biofilm evolution model to select for strains with increased biofilm formation, with and without *maoP*. This enabled us to identify mutations that compensate for the deletion of *maoP* which we hoped would further describe the regulatory involvement of MaoP in biofilm formation. We found that the WT and Δ*maoP* strains evolved along different paths albeit with two identical SNPs being selected in both strains.

Many Δ*maoP* strains recovered from the model did not fully recover to the rdar phenotype and developed SNPs upstream of *fimZ*, the positive regulator of fimbriae, a structure known to be important for adhesion (8,9,11,57). The SNPs were not found, to the best of our knowledge, within regions where regulatory agents bind. Representative strains did not demonstrate increased biofilm formation in pure culture. Instead, such isolates were selected for in the model likely due to their increased adhesion caused by the increased *fimZ* transcript abundance in the cells and hence increased fimbriae production. It is possible that isolates with such SNPs were not recovered from the WT model due to a higher baseline fitness meaning that such mutations provided a smaller relative fitness increase in WT, and correspondingly lower selective pressure for attachment.

### The biofilm evolution model highlights a CsgD-mediated role of MaoP in biofilm formation

All strains that had fully recovered the rdar phenotype in the Δ*maoP* background had one of two SNPs upstream of *csgD*. Despite the absence of *maoP*, representative Δ*maoP* isolates possessing either SNP demonstrated increased biofilm formation, curli, and cellulose production, due to significantly increased *csgD* mRNA levels in the cell in a biofilm.

One SNP was identified within the OmpR binding site, which was also recovered from the WT model. This SNP has been found in clinical isolates with increased biofilm formation (58). OmpR, part of the OmpR-EnvZ two component system to respond to osmolarity, binds in six sites of the *csgD* promoter, this SNP specifically is located within the site where OmpR binding is essential for transcription (43,50,51). The presence of this SNP in Δ*maoP* isolates with increased biofilm formation may suggest an involvement of OmpR in the way in which MaoP regulates biofilm, however the recovery of this SNP also from the WT model and a lack of clear signal suggesting this possibility in the transcriptomic data does not support this hypothesis. The other SNP recovered, only from the Δ*maoP* model, was found to lie in the - 10 binding region for the RNA polymerase. The fact that this was not also recovered from the WT model could suggest a specific interaction of MaoP with this site that should be further investigated.

## Conclusions and future work

In conclusion, we have shown that MaoP regulates biofilm formation via modulating *csgD* mRNA levels in the cell and increasing expression of *csgD* can suppress this deletion phenotype. Whilst it now seems clear that MaoP is needed to support normal levels of *csgD* in the cell, how this interaction is mediated remains unclear. Ongoing work aims to identify the potential binding targets of MaoP that causes the phenotypes seen, including RNA and DNA binding assays to further our understanding of biofilm regulation in *Salmonella*.

## DATA AVAILABILITY

The raw sequencing datasets generated for RNA-seq are available in the NCBI Sequence Read Archive under BioProject accession PRJNA1563442.

## ACKNOWLEDGEMENTS

The authors gratefully acknowledge the Quadram Institute Bioscience sequencing team, David Baker and Beth Baker, for whole genome sequencing. The authors also thank Dr. Katie Noble for her insightful comments on drafts of this paper.

