## Supplementary figures for "*maoP* impacts biofilm formation via *csgD* in *Salmonella* Typhimurium"

**Supplementary material**

**Supplementary figures**

**Supplementary Table 1 Plasmids used in this work**

| Name | Source | Use | Resistance Marker(s) |
| --- | --- | --- | --- |
| pDOC-K-glmS | (1) | Gene doctoring donor plasmid for insertion of DNA constructs as phenotypically neutral Tn7 insertion site (‘glmS site’) in *S.* Typhimurium 14028*S* | Carbenicillin, Kanamycin |
| pDOC-K-glmS-maoP | This project | Insertion of *maoP* for complementation | Carbenicillin, Kanamycin |
| pCP20 | (2) | Removal of antibiotic resistance markers within FLP recombinase recognition sites | Carbenicillin |

**Supplementary Table 2 Primers used in this work**

| Primer name | Sequence (5’-3’) | Use |
| --- | --- | --- |
| STMmaoP-F | GGGCTCGAGACTGTGCTCTCCACAAAATTATTTGTC | Amplify *maoP* with native promoter from *S.* Typhimurium 14028*S*, adding XhoI site at beginning and XmaI site at end for insertion into pDOC-K-glmS |
| STMmaoP-R | AGACCCGGGTTAATCATCAGCTTCGGTGTAGTCTTCTG |  |
| pDOC-F | CATGATTACGCCAAGCTCTAG | Check for insertion of construct into pDOC-K-glmS |
| pDOC-R | GGGTTTTCCCAGTCACGACGT |  |
| STMglmS-F | GGCCAGGGGGATCCACCTGAATAACCTGGG | Check for insertion of DNA construct into chromosome by Gene Doctoring at ‘glmS site’ in *S.* Typhimurium 14028*S* |
| STMglmS-R | ACGTGGCGCTGATTAAAGGCACCGACGTTG |  |
| STMcsgD-upstream-F | GGGGGTCTCTCGCTCTCCACTTTTTTTAATCGCACAT | Amplify promoter region of *csgD* in *S.* Typhimurium for Sanger sequencing |
| STMcsgD-upstream-R | GGGGGTCTCATCGTTTTTTTAAATCTATGCAATAACAGCGAAATG |  |
| STM-RT-csgD-F | CCCTGACGATTATCCCTACCG | RTqPCR for *csgD* in *S.* Typhimurium |
| STM-RT-csgD-R | CTGTAATCCGCTGACCACGT |  |
| STM-RT-gyrB-F | AGGTCTGATTGCGGTGGTTT | RTqPCR for housekeeping gene, *gyrB*, in *S.* Typhimurium |
| STM-RT-gyrB-R | TGCTGTTCTACCGCCGATTT |  |
| STM-RT-fimZ-F | CGCCCGCTCTTATTGCTCTT | RTqPCR for *fimZ* in *S.* Typhimurium |
| STM-RT-fimZ-R | TCTATCCAGGAACATACCCGG |  |

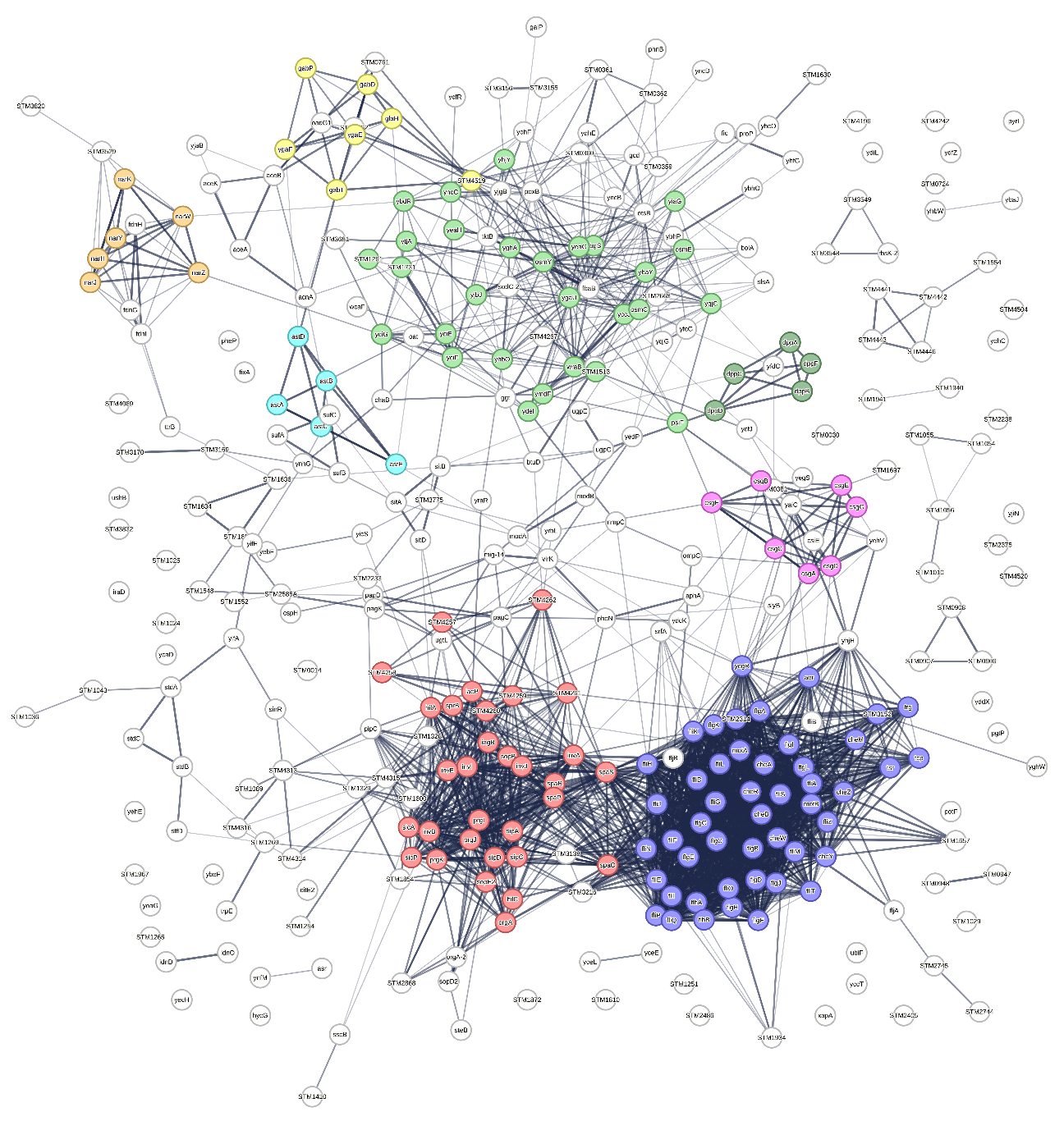

**Supplementary Figure 1 STRING network of differently expressed genes in Δ*maoP* 24-hour biofilm compared to WT**

Default parameters, importing genes from RNA-seq analysis. Coloured to highlight clusters of different cellular processes: blue = CL:5089, Flagellar assembly, and Bacterial chemotaxis ; red = CL:5324, Virulence, and Type I secretion membrane fusion protein, HlyD family ; light green = CL:6425, Mostly uncharacterized, incl. Ferritin-like, and DUF883, N-terminal domain ; pink = CL4538, Mixed, incl. Curlin associated, and Helix_turn_helix, Lux Regulon ; yellow = CL:7828, Succinate-semialdehyde dehydrogenase (NAD+) activity, and Lysine catabolic process ; dark green = CL:6120, Oligopeptide/dipeptide transporter, C-terminal region, and Dipeptide transport ; cyan = CL:2036, arginine catabolic process to succinate ; orange = CL:3949, Nitrate reductase activity.

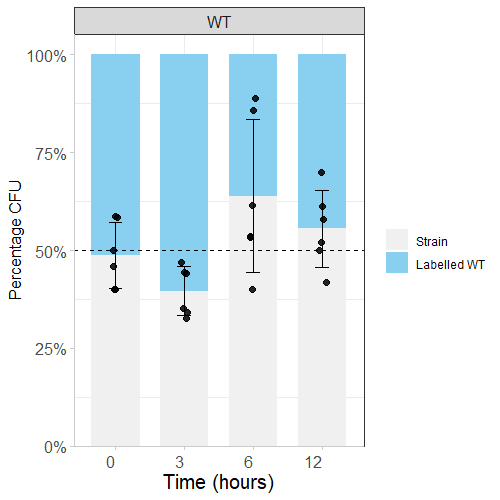

**Supplementary Figure 2 Unlabelled and LacIZ labelled WT do not show significant differences in the bead adhesion model**

Biofilm adhesion competition assay between a LacIZ tagged WT strain and WT. CFU counts taken at Time=0 (planktonic), 3, 6, and 12 hours (biofilm). The mean percentage of mutant (white) and labelled WT (blue) colonies are represented by the grey and blue bars respectively. Error bars show one standard deviation of the percentage of mutant colonies. Points represent six replicates (representing three samples from each of two independent runs completed on different days). One-way ANOVA did not reveal statistically significant differences between timepoints.
